# Pan-Screening the Structural Predictability of Drug Toxicity: Validation of Molecular Fingerprints

**DOI:** 10.64898/2026.09.27.754850

**Authors:** Wei Jiang, Xiang’e Jian, Jie Hu, Mengru Fu, Sufang Xu

## Abstract

**Objective:** To systematically evaluate the predictability of drug chemical structures against the complete set of MedDRA Preferred Terms (PTs), construct a comprehensive structure– adverse reaction landscape, validate the utility of molecular fingerprint representations in pan-screening tasks, and establish quantitative tools for drug safety pre-screening.

**Methods:** Drug–adverse event association data were compiled from SIDER 4.1 and its derivative compilation Nsider, encompassing 2,005 approved drugs, of which 1,120 small-molecule drugs with successfully retrieved chemical structures were retained for analysis. For each Preferred Term (PT) meeting the minimum sample-size threshold (≥25 positive and ≥30 negative drugs; *n* = 768), XGBoost classifiers were trained using MACCS 166-bit fingerprints, with model performance evaluated via 5-fold stratified cross-validation. A PT was designated as high-confidence if it achieved a validation AUC ≥ 0.75, a cross-validation AUC ≥ 0.70, and passed Benjamini–Hochberg false discovery rate (FDR) correction (*q* < 0.05). The entire analytical pipeline was replicated using Morgan ECFP4 2048-bit fingerprints to assess the robustness of fingerprint selection. The robustness of significance screening was independently validated by permutation testing (500 random label permutations), and model generalizability was evaluated through Bemis–Murcko scaffold-based stratified split validation. SHAP (SHapley Additive exPlanations) analysis was employed to identify critical structural features, and mechanism-based case studies further demonstrated concordance between model-identified features and established pharmacophores.

**Results:** Among the 768 modelable PTs, 71 (9.2%) met the high-confidence significance criteria (FDR *q* < 0.05), predominantly comprising nervous system disorders (10 PTs) and psychiatric disorders (9 PTs). The toxicities exhibiting the strongest structural predictability included hyperprolactinaemia (AUC = 0.968), dermatitis acneiform (AUC = 0.966), tardive dyskinesia (AUC = 0.952), and breast tenderness (AUC = 0.979). Under a unified XGBoost hyperparameter configuration, the low-dimensional MACCS fingerprint (166-bit) demonstrated near-equivalent performance to the high-dimensional Morgan fingerprint (2048-bit) in pan-screening tasks (ΔAUC = −0.012, Cohen’s *d* = 0.16), and achieved comparable performance at a substantially lower dimensionality across 18 of 22 organ systems. SHAP analysis identified key structural fragments corresponding to established pharmacophores (six-membered rings, tertiary amine cores, heteroatomic bonds, and bridged heteroatomic chains), and mechanism-based case studies further demonstrated that the model captured structural patterns concordant with known pharmacophores of D2 antagonists, EGFR inhibitors, and related agents. Bemis–Murcko scaffold-based stratified split validation revealed a median AUC decrease of merely 0.042, with 66% of PTs maintaining an AUC ≥ 0.70 under scaffold-split conditions. Immune-mediated and multifactorial endpoints remained structurally opaque under the current data conditions.

**Conclusion:** This study constructed a comprehensive structure–adverse reaction predictability landscape encompassing 768 MedDRA PTs; seventy-one PTs achieved high-confidence criteria. The MACCS 166-bit fingerprint achieved near-equivalent performance to the Morgan 2048-bit fingerprint at a markedly lower dimensionality, and its chemically explicit features enable direct translation of model outputs into structural alerts, providing a computational tool for prospective screening of potential adverse reaction profiles for any novel compound. A tiered assessment strategy is proposed: machine learning ensemble models for high-confidence screening, substructure alert rules for high-specificity exclusion, and pharmacovigilance signal detection for complementary coverage of structurally unpredictable endpoints. Bemis–Murcko scaffold-based stratified split validation further demonstrated that the median generalization performance decrease was merely 0.042 AUC, with 66% of PTs maintaining an AUC ≥ 0.70 under scaffold-split conditions, corroborating the existence of a genuine chemical basis underlying structure–adverse reaction mappings.

## 1. Introduction

Adverse drug reactions (ADRs) account for approximately 5–7% of hospital admissions worldwide and represent one of the leading causes of clinical trial failure and post-marketing drug withdrawal ^1,2^. The capability to predict drug toxicity from chemical structures prior to clinical stages would substantially reduce attrition in drug development and improve patient safety.

Computational toxicology has achieved substantial progress over recent decades. Quantitative structure–activity relationship(QSAR) models, deep neural networks, and ensemble learning methods have been successfully applied to predict specific toxicity endpoints, including cardiotoxicity(hERG inhibition),hepatotoxicity, and respiratory toxicity ^3–6^. However, nearly all published studies have focused on modeling individual predefined endpoints, whereas the pan-predictability of drug structures across thousands of adverse effect endpoints has rarely been systematically explored.

This endpoint-centric paradigm has created knowledge gaps at two levels. First, for pharmaceutical developers evaluating novel chemical entities, the critical question—which adverse effects among thousands of possibilities are structurally predictable for a given compound—cannot be answered through single-endpoint studies, because the absence of systematic predictability assessments and comprehensive landscape construction for the full MedDRA adverse effect terminology from chemical structures precludes medicinal chemists from preemptively identifying which adverse reactions require prioritized attention during the evaluation of new compounds. Second, most toxicity prediction studies lack systematic comparison and empirical justification in the selection of molecular fingerprint types; the choice among MACCS, Morgan, or Klekota–Roth fingerprints is typically not validated against the specific prediction task. A prevalent tendency in molecular representation selection is the default assumption that higher-dimensional fingerprints (e.g., Morgan 2048-bit) afford superior predictive performance by encoding more granular structural information^7,8^, yet this assumption remains to be systematically examined in small-sample toxicity screening tasks. However, the trade-off between fingerprint resolution and statistical power in screening tasks spanning hundreds of endpoints with small positive sample sizes has been rarely discussed in the toxicity screening literature. The crux of this issue is that for small-sample endpoints, although high-dimensional fingerprints encode richer structural information, the concomitant deterioration in the feature-to-sample ratio may offset this information gain.

Pharmacologically, drug toxicities can be broadly classified into three categories: on-target effects, off-target effects, and multifactorial or idiosyncratic reactions. This study hypothesizes that a core subset of drug toxicities is chemically structure-determined. It is further hypothesized that this subset is predominantly driven by well-defined receptor–ligand interactions (on-target and off-target effects), whereas endpoints involving immune activation, metabolic bioactivation, and other polygenic mechanisms may be inherently refractory to prediction from chemical structure alone. If this hypothesis holds, systematic screening would enable the construction of a comprehensive structure–adverse reaction predictability landscape, providing quantitative guidance for medicinal chemistry optimization and risk pre-assessment of novel compounds. This classification furnishes an explanatory framework for the observed heterogeneity in structural predictability in the subsequent analyses.

In this study, we address these gaps through systematic, data-driven screening of all 26,641 MedDRA 27.1 PTs. The study comprises four core components: (1) multi-source SMILES curation from DrugBank, PubChem, NIH CIR, and ChEMBL, with formal classification of unmatched drugs; (2) exhaustive MACCS 166-bit XGBoost modeling and multi-tier significance assessment for PTs meeting the eligibility criteria; (3) replication of the entire analytical pipeline using Morgan ECFP4 2048-bit fingerprints under identical hyperparameters, with empirical comparison to validate fingerprint selection; and (4) SHAP-guided mechanistic case studies to verify whether MACCS bits for three representative drugs correspond to established pharmacophores and clinical toxicity mechanisms. Our objective is to chart the comprehensive landscape through rigorous empirical methods, rather than to construct optimal models for any individual endpoint.

## 2. Methods

The study comprised four key components: (1) integration of drug–adverse event association data from SIDER 4.1 and its derivative Nsider compilation, coupled with chemical structure curation via a four-tier strategy; (2) for each Preferred Term (PT) meeting the minimum sample-size threshold (≥25 positive and ≥30 negative drugs), training of XGBoost classifiers using MACCS molecular fingerprints—the MACCS fingerprint consists of 166 predefined chemical substructure patterns, with each bit corresponding to an explicit chemical feature; (3) replication of the entire analytical pipeline using an independent Morgan circular fingerprint (a 2048-bit representation based on atom neighborhood hash encoding) to empirically validate fingerprint selection; and (4) employment of game-theoretic SHAP (SHapley Additive exPlanations) values to quantify the contribution of each structural feature to model predictions for key substructure identification, followed by validation through pharmacophore case studies.

### 2.1 Data Sources and Preprocessing

Data were integrated from two complementary SIDER 4.1–derived compilations^9^. The standard SIDER 4.1 release contributed 1,558 STITCH compound identifiers, with adverse effect annotations extracted from approved drug labels. The Nsider compilation independently contributed 707 STITCH compound identifiers (non-overlapping with the former), accompanied by OMOP-standardized concept IDs. Merging the two sources yielded 194,846 drug–adverse effect pairs, encompassing 2,005 unique drugs and 4,251 unique adverse effect terms. Adverse effect terms were normalized to MedDRA 27.1 Preferred Terms (PTs) through case-insensitive string matching; all 26,641 PTs served as candidate endpoints, and those matching SIDER annotations upon normalization and meeting the minimum sample-size threshold were retained for modeling. A drug–PT pair was labeled positive (1) if the drug was reported to be associated with that PT in either data source; it was labeled negative (0) if the drug was present in the merged dataset but not associated with that PT.

A four-tier SMILES curation strategy was employed: (1) canonical SMILES for approved small-molecule drugs were retrieved from DrugBank 5.1^10^, with drug name matching performed through case-insensitive exact matching supplemented by 39,348 synonyms; (2) CAS-to-SMILES conversion was conducted via the NIH Chemical Identifier Resolver (CIR); (3) unmatched drugs were queried through PubChem PUG-REST, with automated name standardization (removal of salt-form suffixes, dosage-form descriptors, and parenthetical qualifiers; splitting of combination drugs at semicolons), retaining valid SMILES within a molecular weight range of 50–2000 Da; and (4) as a final recourse, remaining unmatched names were queried via the ChEMBL 34 REST API^11^. Drugs with RDKit-parsable SMILES obtained after cross-referencing were retained for subsequent analysis. Unmatched drugs were systematically classified according to the reason for curation failure.

### 2.2 Molecular Fingerprinting

MACCS 166-bit fingerprints were calculated using RDKit (v2022.9.5) and employed as the primary molecular representation ^12^. The chemical interpretation of each bit was validated individually against the RDKit smartsPatts dictionary (Supplementary Table S3); for example, bit 122 corresponds to tertiary amines, bit 145 to six-membered rings, and bit 130 to heteroatom– heteroatom bonds. This explicit chemical interpretability permits direct translation of SHAP-derived feature importance into chemically comprehensible structural alerts.

To validate the selection of MACCS through empirical comparison, the entire analysis was replicated for all included PTs using Morgan fingerprints (ECFP4, radius = 2, nBits = 2048). All XGBoost hyperparameters were held consistent with the MACCS pipeline to ensure that performance differences were attributable solely to fingerprint type. Statistical comparison between MACCS and Morgan performance was conducted via paired Wilcoxon signed-rank tests on the AUC values of individual PTs. The performance of Morgan fingerprints with independently optimized hyperparameters may differ from the results reported herein. To evaluate the robustness of the hyperparameter configuration, an independent hyperparameter grid search was performed for Morgan fingerprints.

RDKit-generated MACCS fingerprints comprise 167 bits (bits 0–166); bit 0, which is invariantly zero, was excluded, retaining 166 bits as features. Feature names MACCS_n correspond to the native RDKit index n+1 (e.g., MACCS_145 corresponds to RDKit bit 146).

### 2.3 Model Construction and Evaluation

For each PT meeting the minimum sample-size criteria (≥25 positive and ≥30 negative drugs), models were trained using XGBoost^13^with gradient boosted tree configuration (max_depth = 5, learning_rate = 0.05). Class imbalance was addressed by setting scale_pos_weight to the ratio of negative to positive instances (n_negative/n_positive). The dataset was partitioned into 80/20 stratified splits. In 5-fold stratified cross-validation, n_estimators was set to 100 (for validation purposes, resulting in thousands of model fittings across all included PTs), whereas n_estimators = 200 was employed for the final validation model.

A three-tier significance framework was employed for significance assessment; PTs satisfying all three criteria simultaneously were designated as high-confidence predictable PTs: (1) validation AUC ≥ 0.75^14^; (2) 5-fold CV AUC ≥ 0.70; and (3) Benjamini–Hochberg false discovery rate (FDR) correction (*q* < 0.05). As an independent calibration of the primary significance framework, a permutation test–based validation scheme was implemented for each modelable PT (500 random label permutations), with empirical *p*-values calculated and 95% confidence intervals (CIs) for AUC estimated via bootstrap resampling (*n* = 1,000) for each high-confidence predictable PT ^15^. To assess model generalizability, Bemis–Murcko scaffold-based stratified split validation (scaffold split) was performed for all high-confidence PTs: drugs were grouped by Murcko scaffold, and GroupShuffleSplit was applied to ensure that training and test sets contained no shared scaffolds, followed by retraining and evaluation under identical XGBoost hyperparameters.

All analyses were performed in Python 3.11.7, using scikit-learn 1.2.2, XGBoost 2.1.4, SHAP 0.49.1, and RDKit 2022.9.5.

### 2.4 SHAP Interpretability Analysis

For each high-confidence predictable PT, SHAP values were computed for the final XGBoost model using SHAP TreeExplainer ^16^, with a background dataset of *n* = 80 and an explanation dataset of *n* = 50. The top-5 MACCS bits were reported for each PT (ranked by mean absolute SHAP value). All SHAP-identified MACCS bits were cross-referenced against RDKit smartsPatts definitions to ensure accurate chemical interpretation.

To further validate whether SHAP-identified features capture authentic pharmacophore information, mechanism-based case studies were performed on three representative drugs: (1) haloperidol (D2 receptor antagonist), corresponding to tardive dyskinesia and hyperprolactinaemia; (2) gefitinib (EGFR tyrosine kinase inhibitor), corresponding to dermatitis acneiform; and (3) morphine (μ-opioid receptor agonist), corresponding to hypoventilation. For each drug, analyses were conducted according to the following procedure: first, MACCS bits triggered by the drug were computed via RDKit; subsequently, those bits appearing in the SHAP top-5 for the relevant toxicities were translated into specific chemical groups through SMARTS definitions; finally, these groups were evaluated against published pharmacophore–target association knowledge to determine whether they were concordant with established pharmacological mechanisms.

## 3. Results

### 3.1 Pan-Screening

The multi-source SMILES curation pipeline retrieved valid chemical structures for 1,120 of the 2,005 SIDER-annotated drugs (55.9% coverage rate). Among the remaining 885 unmatched drugs, 631 (71.3%) were combination drug entries, and 237 lacked structures in all queried databases. Among the 26,641 MedDRA PTs, 3,903 (14.6%) had at least one matching annotation, 768 met the minimum modeling threshold, and 71 satisfied all high-confidence criteria (FDR q < 0.05).

This number increased to 78 following permutation test calibration. High-confidence predictable PTs accounted for 9.2% of modelable PTs and 0.27% of the full MedDRA PT set. In other words, screening across all 26,641 PTs indicated that, under current data scales and modeling conditions, only a minority of adverse effect endpoints can be predicted with relatively high accuracy from chemical structure given the available data. The validation AUC distribution for the 768 modelable PTs exhibited a median of 0.637 (interquartile range [IQR], 0.573–0.698; Figure 1).

**Figure 1.**
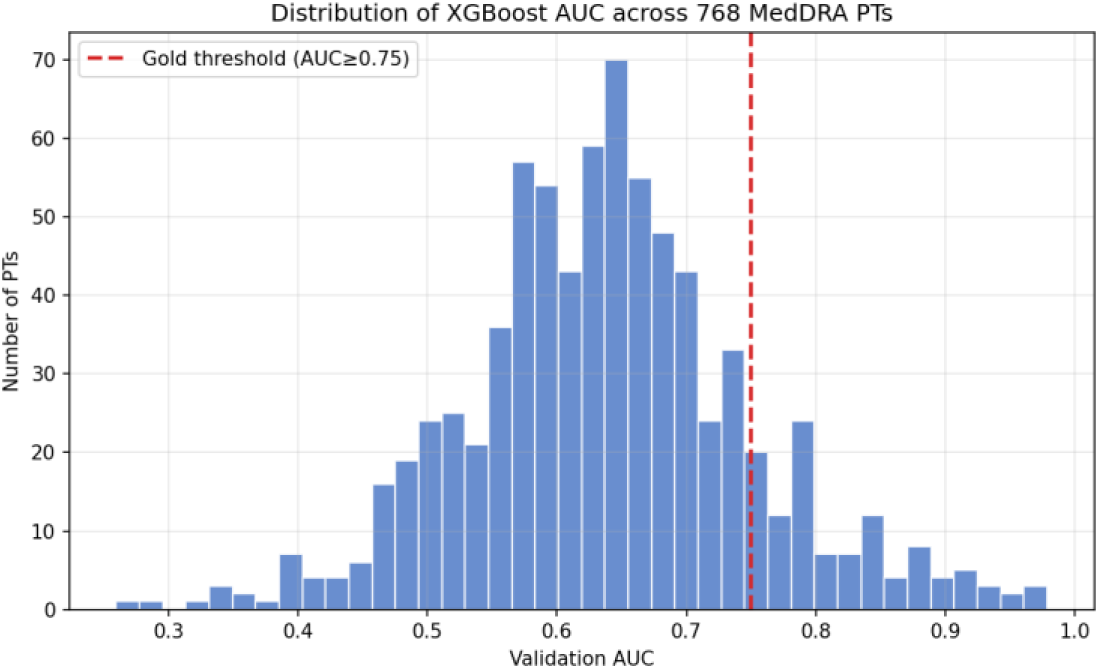
Histogram of the validation AUC distribution for the 768 modelable PTs. The dashed line denotes the high-confidence threshold (AUC = 0.75). The distribution is approximately symmetric (median = 0.637), with the right tail extending to 0.979, representing the minority of structurally predictable endpoints.

The bulk of the distribution is approximately symmetric about the median; however, the right tail extends to 0.979, signifying that a minority of PTs attained high structural predictability under the present modeling conditions.

### 3.2 Distribution by System Organ Class

The 71 high-confidence predictable PTs spanned 20 of 27 System Organ Classes (SOCs) and were significantly enriched in nervous system disorders and psychiatric disorders (Figure 2).

**Figure 2.**
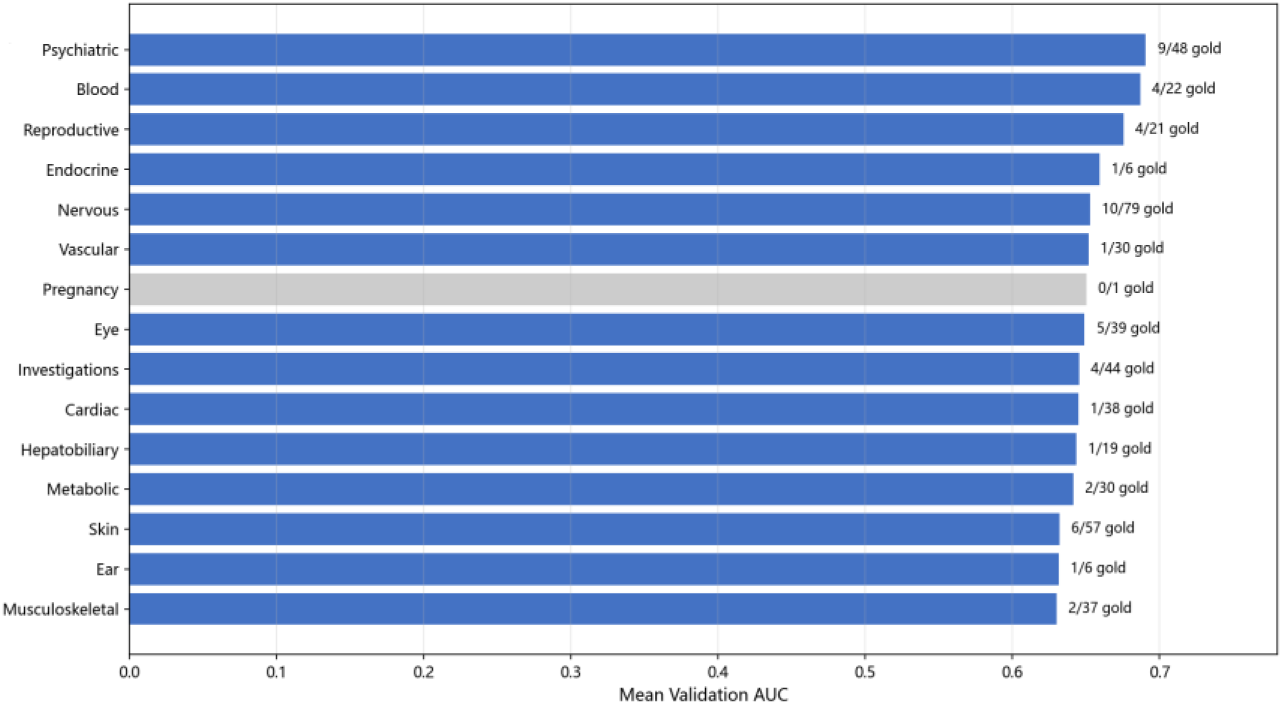
Distribution of high-confidence predictable PTs across SOCs (top 15).

Nervous system disorders contributed 10 high-confidence predictable PTs, whereas psychiatric disorders contributed 9. Other enriched SOCs encompassed skin disorders (7 PTs), general disorders (6 PTs), eye disorders (5 PTs), and blood/lymphatic system disorders (4 PTs). The respiratory system SOC yielded 4 high-confidence predictable PTs.

### 3.3 Ranking of Structural Predictability

Table 1 lists the top ten high-confidence predictable PTs ranked by validation AUC (Figure 3). Breast tenderness (AUC = 0.979), hyperprolactinaemia (0.968), dermatitis acneiform (0.966), and tardive dyskinesia (0.952) ranked highest. All top 10 PTs achieved AUCs exceeding 0.91, suggesting the existence of a toxicity subset amenable to highly accurate structure-based prediction. The key SHAP MACCS bits for each PT were cross validated against SMARTS definitions, pointing to specific chemical determinants.

**Figure 3.**
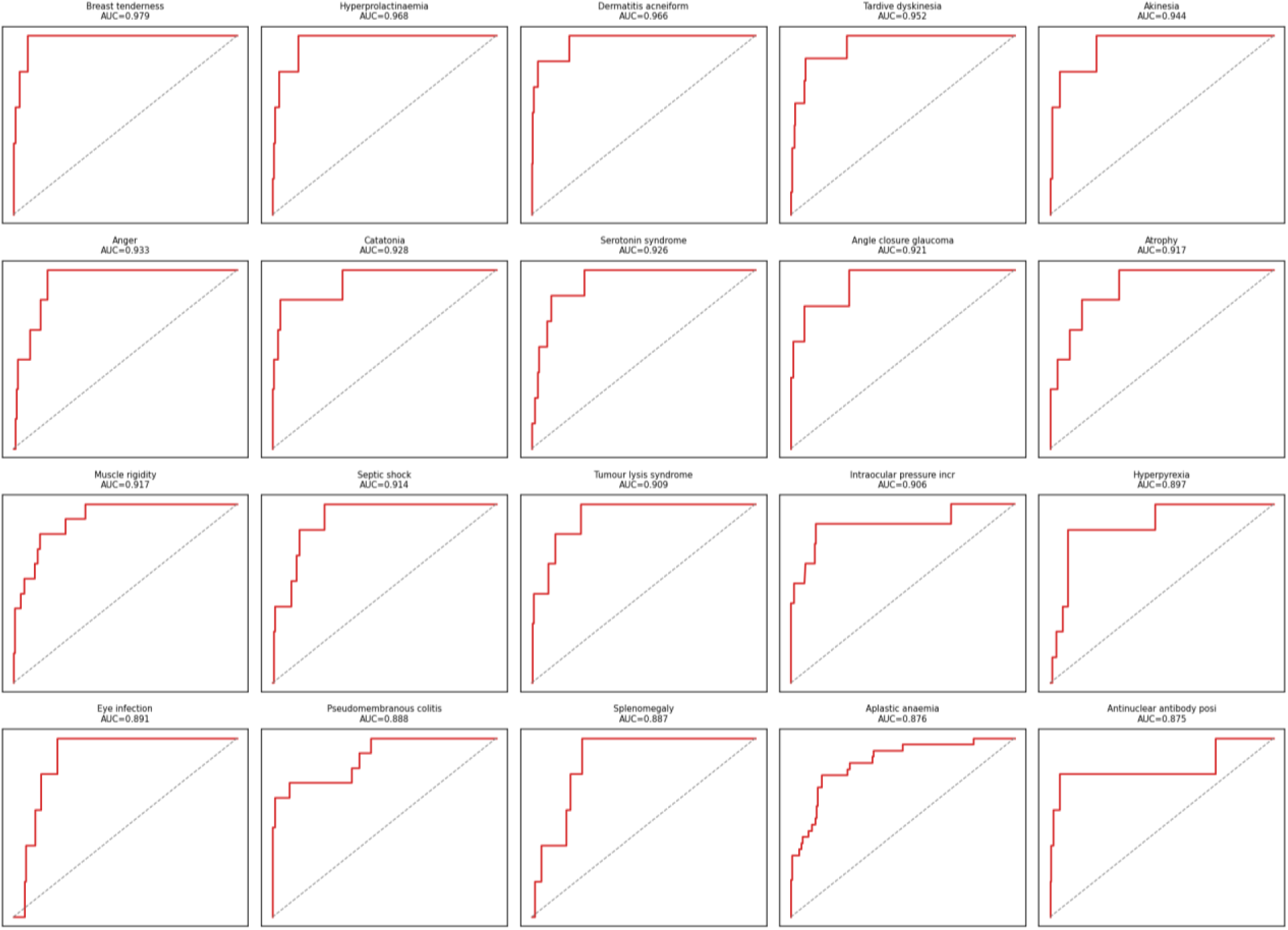
Receiver operating characteristic (ROC) curves of the top 15 models ranked by validation AUC.

**table 1.** Predictive performance of models for drug-induced adverse events and key MACCS structural fragments identified by SHAP.

| PT | SOC | AUC | CV AUC | F1 | n+ | Key SHAP MACCS bits |
| --- | --- | --- | --- | --- | --- | --- |
| Breast tenderness | Reproductive system and breast disorders | 0.979 | 0.822 | 0.667 | 27 | 92; 145; 81 |
| Hyperprolactinaemia | Endocrine disorders | 0.968 | 0.852 | 0.600 | 25 | 85; 131; 136 |
| Dermatitis acneiform | Skin and subcutaneous tissue disorders | 0.966 | 0.860 | 0.714 | 36 | 148; 71; 110 |
| Tardive dyskinesia | Nervous system disorders | 0.952 | 0.873 | 0.526 | 38 | 145; 149; 130 |
| Akinesia | Nervous system disorders | 0.944 | 0.804 | 0.667 | 27 | 130; 89; 139 |
| Anger | Psychiatric disorders | 0.934 | 0.730 | 0.500 | 31 | 130; 132; 95 |
| Catatonia | Psychiatric disorders | 0.928 | 0.854 | 0.600 | 30 | 154; 79; 89 |
| Serotonin syndrome | Nervous system disorders | 0.926 | 0.912 | 0.421 | 35 | 145; 85; 149 |
| Angle closure glaucoma | Eye disorders | 0.922 | 0.856 | 0.571 | 26 | 135; 141; 53 |
| Atrophy | General disorders and administration site conditions | 0.917 | 0.857 | 0.500 | 30 | 138; 119; 145 |

To assess the stability of AUC point estimates for high-confidence predictable PTs, 95% confidence intervals (CIs) for AUC were calculated for each high-confidence predictable PT via bootstrap resampling (*n* = 1,000). The mean CI width for the 71 high-confidence predictable PTs was 0.273 (median, 0.238). Stratification by CI width yielded a high-confidence stratum (CI < 0.25; 43 PTs; mean AUC, 0.856), a medium-confidence stratum (CI 0.25–0.50; 29 PTs; mean AUC, 0.807), and a low-confidence stratum (CI > 0.50; 6 PTs; mean AUC, 0.784). The narrowest CIs were observed for breast tenderness ([0.937, 0.999]) and dermatitis acneiform ([0.926, 1.000]), whereas the widest were observed for telangiectasia ([0.320, 1.000]) and subcapsular cataract ([0.336, 1.000]), indicating that assessments of structural predictability for small-sample endpoints should be interpreted cautiously in conjunction with CI width.

### 3.4 Comparison of MACCS and Morgan Fingerprints

Replication of the full 768-PT analysis using Morgan ECFP4 2048-bit fingerprints yielded three key findings (Figure 4).

**Figure 4.**
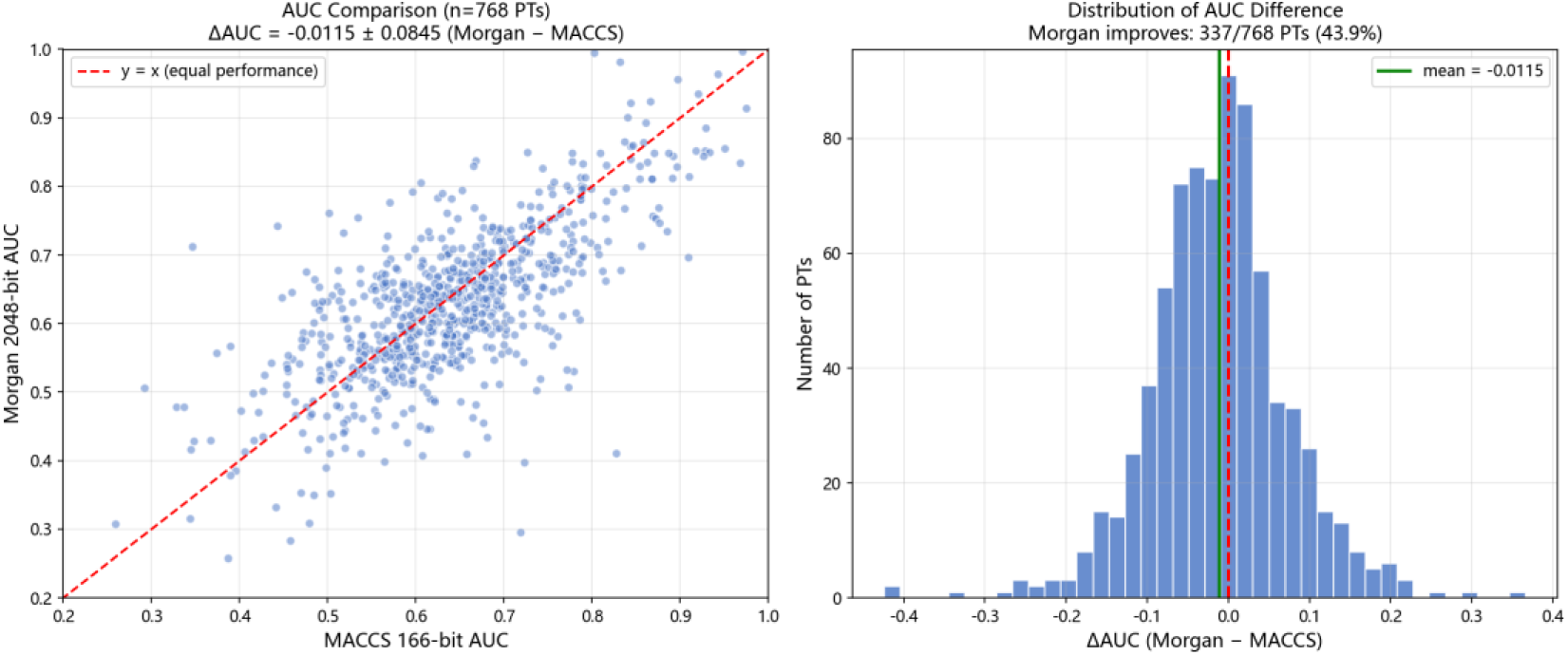
Comparative performance of Morgan ECFP4 2048-bit versus MACCS 166-bit fingerprints. (a) Scatter plot of AUC values across 768 PTs (diagonal line denotes equal performance). (b) Histogram of ΔAUC (Morgan − MACCS) distribution, with the red dashed line indicating Δ = 0.

Overall performance. Under a unified hyperparameter configuration, the mean AUC across 768 PTs was 0.639 for MACCS versus 0.627 for Morgan (ΔAUC = −0.012, 95% confidence interval [CI] [−0.017, −0.007], paired Wilcoxon signed-rank test, *p* = 3.5 × 10⁻⁵). The standardized effect size (Cohen’s *d*) was 0.16, falling below the conventional small-effect threshold (*d* = 0.2), indicating a negligible practical difference in performance between the two fingerprints under the current configuration. MACCS yielded a higher AUC in 429 PTs (55.9%), Morgan in 337 PTs (43.9%), with 2 PTs resulting in a tie. The number of high-confidence predictable PTs (FDR *q* < 0.05) under a unified z-score threshold was 71 for MACCS versus 58 for Morgan (Δ = −13); this increased to 78 for MACCS following permutation test calibration.

Stratification by System Organ Class. Among the 22 SOCs containing at least 5 PTs, MACCS achieved a higher mean AUC in 18. The systems exhibiting the greatest numerical difference favoring MACCS were psychiatric disorders (ΔAUC = −0.037), blood and lymphatic system disorders (−0.035), and endocrine disorders (−0.032). Morgan yielded a higher mean AUC in only 4 SOCs (≥5 PTs): injury, poisoning and procedural complications (+0.052), immune system disorders (+0.021), eye disorders (+0.019), and renal and urinary disorders (+0.008).

Under the current unified hyperparameter modeling configuration, the MACCS 166-bit fingerprint exhibited a statistically significant yet small-effect difference in this pan-screening task (Cohen’s d = 0.16, 13 additional high-confidence predictable PTs, and a higher mean AUC in 18 of 22 SOCs at the organ-system level).

### 3.5 Chemical Basis of Predictability

SHAP feature importance analysis identified four chemical patterns recurrently observed across the 71 high-confidence predictable PTs (Figure 5). The most frequently occurring bits included hydrogen-bearing heteroatoms (MACCS_130, 15/71 PTs), bridged heteroatomic chains (MACCS_53, 14/71 PTs), and oxygen atoms (MACCS_145, 14/71 PTs; MACCS_139, 9/71 PTs). The SHAP feature combinations of individual PTs exhibited differential distributions at the SOC level: SHAP features for nervous system disorders were predominantly carbon–heteroatom skeletons (MACCS_85) and bridged heteroatomic chains (MACCS_53, MACCS_79), whereas skin disorders were characterized by high-frequency combinations of oxygen-related features and halogenated aromatic rings (MACCS_86).

**Figure 5.**
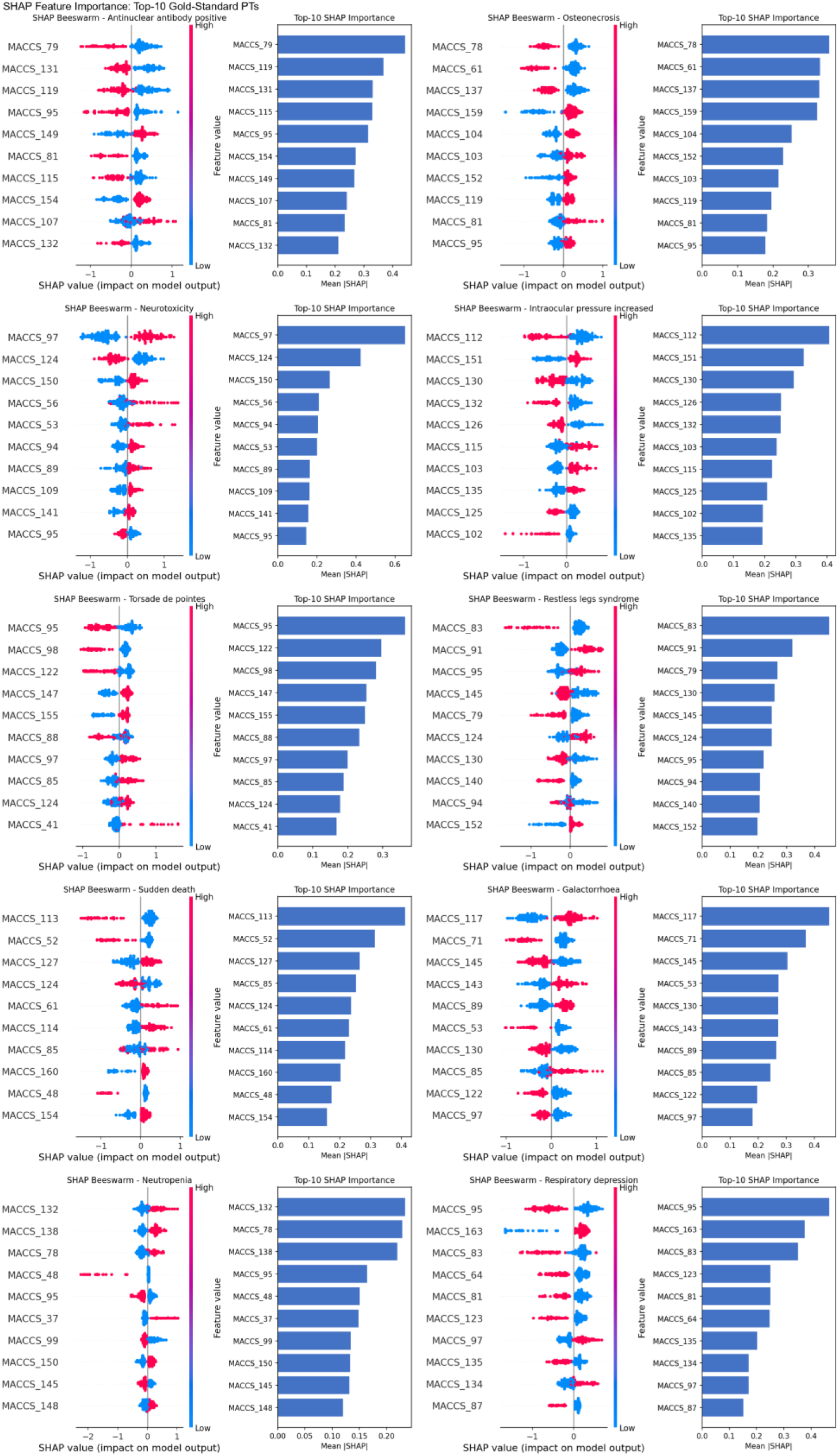
Heatmap of SHAP features importance for the 71 high-confidence predictable PTs. Each row corresponds to an individual PT, each column to a MACCS fingerprint bit, and color intensity encodes the mean absolute SHAP value. System Organ Class (SOC) affiliations are annotated on the right.

### 3.6 Significance Calibration and Generalization Validation

As an independent calibration of the z-score significance framework, permutation testing (500 random label permutations) calibrated the number of high-confidence predictable PTs from 71 to 78, demonstrating high concordance between the two approaches (overlap, 69/71 = 97.2%) and validating the robustness of significance screening. Stratification of bootstrap 95% confidence intervals (CIs) showed that, among the 78 high-confidence predictable PTs, 43 exhibited CI widths <0.25, 29 exhibited widths of 0.25–0.50, and 6 exhibited widths >0.50 (mean CI width, 0.273). The upper bounds of CIs approaching 1.000 for multiple PTs were attributable to the combined effect of the AUC upper-bound constraint and small sample sizes; accordingly, the CI lower bound and width should be regarded as the principal evaluation criteria.

Bemis–Murcko scaffold-based stratified split validation demonstrated that the 71 high-confidence predictable PTs achieved a mean AUC of 0.739 under scaffold-split conditions (0.773 under random split), with a median ΔAUC of −0.042; 47 PTs (66%) maintained an AUC ≥ 0.70 under scaffold-split, and 60 PTs (85%) maintained an AUC ≥ 0.65, indicating a slight optimistic bias (approximately 0.03–0.04 AUC) in performance estimates under random partitioning, yet confirming that the structural signal overall passed the more stringent scaffold generalization test.

## 4 Discussion

### 4.1 Empirical Comparison of Fingerprint Selection

Regarding fingerprint representations, the low-dimensional MACCS 166-bit fingerprint achieved predictive performance approximately equivalent to that of the high-dimensional Morgan 2048-bit fingerprint (ΔAUC = −0.012, Cohen’s d = 0.16), despite utilizing a feature count far below that of the latter. The statistical significance of the observed difference stemmed from the high statistical power afforded by the large-sample paired test and should not be overinterpreted as representing a substantively meaningful performance advantage.

Independent hyperparameter grid search for Morgan fingerprints on representative PTs further substantiated the methodological rationale underlying the unified hyperparameter design. Taking interstitial lung disease (ILD; n = 1,060, 82 positives) as an example, 5-fold cross-validation across 96 hyperparameter combinations (max_depth 3–8, learning_rate 0.01–0.2, n_estimators 100–500) revealed that the optimal cross-validation configuration (max_depth = 8, learning_rate = 0.2, n_estimators = 100) achieved a CV AUC of 0.710; however, the AUC on the independent test set reserved through an 80/20 split was merely 0.588, inferior to the test-set AUC of 0.599 obtained with the default configuration (max_depth = 6, learning_rate = 0.05, n_estimators = 200). These findings indicate that, for toxicity screening tasks with limited positive sample sizes (n⁺ ≈ 25–963), the optimal cross-validation hyperparameters for Morgan fingerprints are susceptible to pronounced overfitting, rendering the “optimal performance” derived therefrom unreliable.

The near-equivalent performance achieved by MACCS at a substantially lower dimensionality than Morgan may be attributed to two factors. First, the 166 chemically explicit features of MACCS exhibit high information density, with each bit directly corresponding to an interpretable chemical substructure, thereby enabling direct translation of SHAP-derived importance into actionable chemical alerts. Second, under conditions of limited positive samples (median n⁺ = 77), the reduced dimensionality mitigates the risk of overfitting. The modest numerical differences favoring Morgan in four SOCs (injury, poisoning and procedural complications; immune system disorders; eye disorders; and renal and urinary disorders) suggests that multi-pathway toxicities may benefit from the additional information capacity of high-dimensional representations; nevertheless, this conjecture awaits empirical validation.

### 4.2 Pharmacostructural Validation of SHAP Features

The intended objective of SHAP analysis is to identify the combinations of MACCS bits most pertinent to the prediction of specific toxicities. Ideally, an individual structural motif might exhibit a one-to-one correspondence with a particular toxicity; however, the present study found that no single high-importance bit identified by SHAP could independently account for the prediction of any given toxicity. Instead, the predictive determination for each toxicity arose from the concerted contribution of an ensemble of bits (Figure 6).

**Figure 6.**
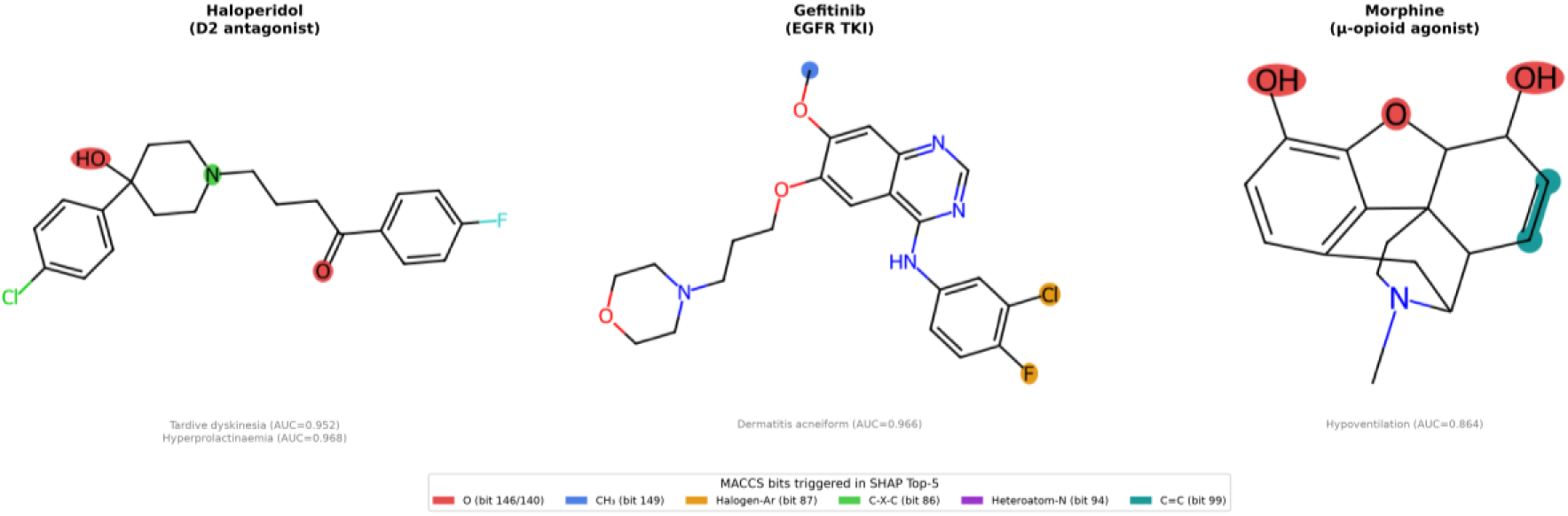
Comparison of chemical structures and top-5 SHAP MACCS bits for three representative drugs.

The SHAP top 5 for tardive dyskinesia (AUC = 0.952) comprised MACCS_145 (oxygen atoms), MACCS_149 (ring patterns), MACCS_130 (hydrogen-bearing heteroatoms), MACCS_148 (methyl groups), and MACCS_101 (heteroatom–oxygen bonds), with hydrogen-bearing heteroatoms and ring patterns as the dominant features. Haloperidol triggered two of these bits (MACCS_145 and MACCS_148). For hyperprolactinaemia (AUC = 0.968),another on-target toxicity of this drug, the SHAP top-5 consisted of MACCS_85 (C–X–C skeleton), MACCS_131 (O–CH₂–C linkage), MACCS_136 (non-carbon atoms within rings), MACCS_106 (halogen branches), and MACCS_148 (methyl groups), dominated by carbon– heteroatom skeletons and oxygen-bridge linkages, forming a distinctive feature combination from that of tardive dyskinesia. Haloperidol likewise triggered two bits in this profile (MACCS_85 and MACCS_148). Although the two SHAP combinations shared only MACCS_148 (methyl groups) in common, both are consistent with the shared chemotypic features of butyrophenone- and piperidine-class D2 antagonists.

The SHAP top 5 for dermatitis acneiform (AUC = 0.966) included MACCS_148 (methyl groups), MACCS_71 (O–C–C–O), MACCS_86 (halogen–aromatic rings), and MACCS_104 (branched rings); gefitinib triggered MACCS_148 and MACCS_86. The halogenated aromatic ring corresponding to MACCS_86 represents a shared structural fragment of quinazoline-class EGFR inhibitors. This combination is characterized by halogenated aromatic rings and oxygen-bridge linkages, markedly distinct from the ring-pattern/hydrogen-bearing-heteroatom combinations associated with neuroleptic-class toxicities. The SHAP top 5 for hypoventilation (AUC = 0.864) comprised MACCS_135 (carbonyl groups), MACCS_93 (heteroatom–nitrogen bonds), MACCS_139 (oxygen atoms), and MACCS_98 (carbon–carbon double bonds); morphine triggered the first three, which collectively encompass the phenolic hydroxyl, ether oxygen, and tertiary amine features of the morphinan skeleton.

The SHAP top 5 combination patterns of the three drug classes were mutually distinguishable at the level of dominant feature categories: neuroleptic-class toxicities were characterized by high-frequency combinations of ring patterns and hydrogen-bearing heteroatoms, EGFR inhibitors by halogenated aromatic rings and oxygen-bridge linkages, and opioids by carbonyl groups and heteroatom–nitrogen bonds. The model automatically clustered chemotype-related structural profiles by combining multiple elementary chemical descriptors, without manual input of prior knowledge; partial overlap between these combinations and established pharmacophores provides indirect concordance evidence indicating that predictions for high-confidence PTs are consistent with known pharmacological knowledge.

### 4.3 Practical Implications

These findings provide a preliminary framework for a practical tiered assessment approach, the PT-level mapping and AUC threshold boundaries between tiers of which require further calibration on independent external validation data. Tier 1: For high-confidence predictable PTs with high AUC (≥0.75), MACCS combined with XGBoost ensemble models are employed for high-confidence screening, with high-probability predictions prioritized for review. Tier 2: For toxicities with intermediate AUC (0.65–0.75), the chemically explicit features of MACCS are leveraged to construct substructure-based structural alert rules, enabling high-specificity exclusion. Tier 3: For endpoints with limited model predictive capacity (AUC < 0.65), traditional pharmacovigilance methods (e.g., disproportionality analysis in FAERS and VigiBase, literature signal detection, and post-marketing cohort studies) serve as the primary assessment modality.

An interactive online platform integrating results for all 768 modelable PTs supports compound structure–based queries of predicted adverse reaction profiles across high-confidence PTs, enabling immediate application of this tiered framework to early safety assessment of new chemical entities.

### 4.4 Related Studies

In the field of computational toxicology, numerous studies have demonstrated the predictive value of structural features for organ-specific toxicities. Li et al. (2025) reported AUCs >0.85 on eight respiratory toxicity endpoints using deep neural networks combined with 4860-bit Klekota–Roth fingerprints; Lee and Yoo (2024) developed the InterDILI model, which achieved interpretable drug-induced liver injury (DILI) prediction by integrating permutation feature importance with attention mechanisms ^17^; Wu et al. (2023) proposed GeoDILI based on geometric deep learning ^18^; Seal et al. (2024) proposed DILIPredictor, which provided MACCS substructure-based interpretation ^19^; Shi et al. (2022) and Gong et al. (2022) established a consensus model and a multi-fingerprint ensemble approach, respectively, for nephrotoxicity prediction ^15, 20^; and Liu et al. (2025) conducted systematic cardiotoxicity prediction using deep learning methods ^21^. These studies have demonstrated structural predictive capacity within specific organ systems; however, all adopted single-system or single-endpoint designs, and none performed head-to-head empirical comparisons of molecular fingerprint selection.

Compared with the aforementioned studies, the present study offers a complementary perspective along three dimensions. First, it constructed a cross-organ-system adverse reaction predictability landscape from chemical structure, rather than pursuing model optimization for individual endpoints. This landscape encompasses 768 MedDRA PTs meeting the minimum modeling threshold, of which 71 achieved high confidence under stringent statistical criteria (FDR *q* < 0.05). Second, through empirical comparison across the full PT space, it validated the effectiveness of low-dimensional chemically explicit fingerprints in pan-screening tasks, consistent with the independent findings of Orosz et al. (2022) and Kim et al. (2024) regarding the interpretability and robustness of MACCS in single-endpoint ADME–Tox tasks, thereby providing cross-endpoint empirical evidence for fingerprint selection. Third, by tracing feature importance to specific chemical groups through SHAP analysis and benchmarking against established pharmacophore knowledge, it furnishes a methodological framework for the pharmacological interpretation of such models.

### 4.5 Limitations

This study has several limitations that should be considered when interpreting the results. First, drug coverage: the 55.9% SMILES matching rate was constrained by the inability to represent combination drugs in the SIDER data—which accounted for 71.3% of unmatched drugs—as single molecular entities. The excluded combination drugs may have contributed disproportionately to the annotations of specific PTs, and their exclusion may introduce coverage bias in certain PTs. Second, label quality and binary simplification: SIDER annotations reflect adverse effects reported in drug labels rather than causally confirmed toxicological relationships; using label-based annotations as training targets may render the predictive performance of some high-AUC PTs susceptible to systematic annotation bias at the drug-class level. The permutation testing framework and bootstrap CIs partially mitigate overfitting risk but cannot correct systematic annotation bias. Stratified modeling using SIDER frequency annotations and drug-class exclusion sensitivity analyses represent directions for subsequent improvement. Third, discrimination between statistical power and intrinsic unpredictability: among the 768 modelable PTs, only 78 (10.2%) met the permutation test–calibrated high-confidence criteria; this study cannot distinguish whether the remaining PTs are intrinsically insensitive to chemical structure or whether their sample sizes were insufficient to detect structural signals. Under the constraint of a median of merely 77 positive drugs, modest structure–toxicity effects may remain undetectable. Accordingly, the label “structurally unpredictable” should be interpreted as “no predictable signal was detected under the current data and threshold conditions.” Stratification of bootstrap 95% CIs further supports this cautious interpretation: among the 78 high-confidence predictable PTs, only 43 (55%) exhibited CI widths <0.25, whereas 6 PTs exhibited CI widths >0.50 (e.g., telangiectasia, CI = [0.320, 1.000]), indicating that the AUC point estimates for some high-confidence predictable PTs are themselves subject to considerable uncertainty. Fourth, external validation: current model evaluation relies on internal random partitioning and scaffold-based stratified split validation; validation against independent pharmacovigilance databases remains to be completed in future work.

## 5 Conclusion

This study performed systematic, data-driven screening of structural predictability across the complete set of 26,641 MedDRA Preferred Terms. Among PTs satisfying the modeling threshold, 78 met high-confidence criteria following permutation test calibration, with nervous system disorders and psychiatric disorders as the predominant categories. For PTs that did not meet these criteria, some may not be inherently refractory to structure-based prediction but rather limited by insufficient sample sizes under the current data constraints.

Under a unified XGBoost hyperparameter configuration, the MACCS 166-bit fingerprint exhibited predictive performance approximately equivalent to that of the Morgan 2048-bit fingerprint, with a statistically significant yet small-effect difference (ΔAUC = −0.012, 95% confidence interval [CI] [−0.017, −0.007], p = 3.5 × 10⁻⁵, Cohen’s d = 0.16). Under a unified z-score threshold, MACCS identified 13 additional high-confidence predictable PTs relative to Morgan (71 versus 58), which further increased to 78 for MACCS following permutation test calibration. Independent hyperparameter grid search for Morgan on representative PTs revealed pronounced overfitting of the optimal cross-validation hyperparameters, further substantiating the methodological rationale underlying the unified hyperparameter design.

SHAP analysis identified high-contribution MACCS features as fundamental chemical descriptors prevalent in drug molecules (hydrogen-bearing heteroatoms, bridged heteroatomic chains, oxygen atoms, carbon–heteroatom skeletons, and halogenated aromatic rings); case validation demonstrated concordance between these features and the authentic chemical groups present in drug molecules. Predictable toxicities were concentrated in PTs associated with target-specific pharmacological mechanisms, whereas immune-mediated and multifactorial endpoints remained structurally opaque.

To facilitate practical application of the research findings, this study developed an accompanying interactive online platform (https://landscape.drugpred.com/), which is freely accessible to the academic community without registration. The platform offers two core functionalities. The first is a landscape query module enabling interactive browsing of structural predictability modeling results for all 768 PTs, with validation AUC, cross-validation AUC, permutation test p-values, and bootstrap 95% confidence intervals available for online inspection for each PT. The second is an online prediction module: upon user input of the target compound’s SMILES string, the platform instantaneously returns its adverse reaction prediction profile across the 78 permutation test–calibrated high-confidence PTs, annotating the key MACCS structural bits underpinning each prediction with SHAP values; the corresponding chemical groups are simultaneously highlighted on the molecular structure, enabling medicinal chemists to rapidly pinpoint potential structural alerts. The complete 768-PT modeling dataset, together with detailed performance metrics for the 71 high-confidence PTs (FDR q < 0.05) and the 78 calibrated PTs, is publicly available on the platform, enabling researchers to conduct adverse reaction profile pre-screening for novel compounds and to extend external validation.

## Ethics declarations

## Ethics approval and consent to participate

Not applicable.

## Consent for publication

Not applicable.

